# Predictive coding over the lifespan: Increased reliance on perceptual priors in older adults – a magnetoencephalography and dynamic causal modelling study

**DOI:** 10.1101/178095

**Authors:** Jason S. Chan, Michael Wibral, Patricia Wollstadt, Cerisa Stawowsky, Mareike Brandl, Saskia Helbling, Marcus Naumer, Jochen Kaiser

## Abstract

Aging is accompanied by unisensory decline; but to compensate for this, two complementary strategies are potentially relied upon increasingly: first, older adults integrate more information from different sensory organs. Second, according to predictive coding (PC) we form ‘templates’ (internal models or ‘priors’) of the environment through our experiences. It is through increased life experience that older adults may rely more on these templates compared to younger adults. Multisensory integration and predictive coding would be effective strategies for the perception of near-threshold stimuli, but they come at the cost of integrating irrelevant information. Their role can be studied in multisensory illusions because these require the integration of different sensory information, as well as an internal model of the world that can take precedence over sensory input. Here, we elicited a classic multisensory illusion, the sound-induced flash illusion, in younger (mean: 27 yrs) and older (mean: 67 yrs) adult participants while recording the magnetoencephalogram. Older adults perceived more illusions than younger adults. Older adults had increased pre-stimulus beta(β)-band activity compared to younger adults as predicted by microcircuit theories of predictive coding, which suggest priors and predictions are linked to β-band activity. In line with our hypothesis, transfer entropy analysis and dynamic causal models of pre-stimulus MEG data revealed a stronger illusion-related modulation of cross-modal connectivity from auditory to visual cortices in older compared to younger adults. We interpret this as the neural correlate of increased reliance on a cross-modal predictive template in older adults that is leading to the illusory percept.

## Introduction

Predictive coding theory suggests that our perceptual experience is determined by a fine balance between internal predictions based on priors acquired over the course of our lives and incoming sensory evidence. Sensory evidence and priors are thought to be fused in a Bayesian way, to arrive at a posterior representing the best guess regarding the state of the world, producing our perception. Aging research offers an opportunity to probe this suggestion, as the amount of information accumulated in our priors increases throughout one’s lifetime, while the precision of unisensory evidence degrades at later stages in life. The first factor will strengthen the influence of predictions, while the second reduces the influence of unisensory evidence. Together, they should tip the balance to a state, where perception is increasingly dominated by predictions. Investigating this change in balance is possible using perceptual illusions that arise when predictions take precedence over sensory evidence, such as the sound-induced flash illusion (SiFi).

The SiFi is the perception of two visual flashes when only one flash is indeed presented along with two auditory beeps, in relatively short succession. This illusion occurs because the auditory modality has a higher temporal acuity compared to the visual system^1-3^, thus we instinctively rely on cross-modal predictions generated by the auditory system^4^. Accordingly, older adults perceive more SiFi across a wider temporal binding window (TBW) compared to young adults^1-5^. This is because older adults are more likely to integrate multisensory stimuli compared to young adults^6^. This was originally thought of as beneficial, as it effectively compensated for the “loss of sensitivity” in unisensory acuity^7^. However, it can also have a detrimental effect, as older adults integrate more sensory information across a longer time span, irrespective of its relevance to a task, compared to younger adults^4^.

To date, the neural underpinnings as to how older adults integrate more information over time are far from understood. However, following the line of argument of predictive coding, it is possible that older adults rely on increased top-down ‘template’ information (i.e. priors) compared to young adults. Perceptual illusions rely on such perceptual ‘templates’ of our environment, which are ultimately violated, but still take precedence. The internal ‘template’ relevant/responsible for the SiFi is that an auditory event should be accompanied by a visual object – thus, when one flash is presented accompanied by two beeps, an additional visual object is perceived^8,9^.

If the predictive coding account is correct in its relation to aging and multisensory integration, then (i) illusions based on predictions should become more frequent with age. (ii) The neurophysiological signature of the illusory perception should also be found as a general marker when comparing young versus older participants. (iii) Due to the predictive nature, this signature should be found preceding the illusion, and (iv) it should be found in the β-band -according to recent neurophysiological accounts of predictive coding^10^. and results from our group^11^. (v) The neurophysiological correlates of the aging process should manifest as network effects in terms of information transfer and effective connectivity, where brain areas generating reliable predictions should increase their influence over other brain areas that deliver less precise sensory evidence.

To assess the neurophysiological correlates of illusory perception and effects of aging, we used magnetoencephalography (MEG) combined with beamformer source reconstruction. For the analysis of network effects, we used a novel combination of information theory, in particular transfer entropy (TE) estimation, and dynamic causal modelling (DCM). TE and DCM are complementary techniques; TE quantifies information transfer between network nodes, i.e. it focuses on network links that channel new information into a node for computation^12^. In terms of inference, TE is an exploratory technique while DCM models the physiological coupling between hidden states of the network nodes^13^, and is a confirmatory approach based on comparing models. Hence, we used the TE-derived network as the basis for a family of DCM models to confirm the structure of the network relevant for the perception of the SiFi, through model comparison. In the winning network, we then looked for quantitative variations in connection strength with age indicative of increased influences of cross-modal predictions in older participants.

## Results

### Increased SiFi in Older Participants

In order to determine the earliest illusion stimulus-onset asynchrony (SOA) which will differentiate the age groups, participants were presented with a version of the SiFi to assess their temporal binding window outside the MEG. In the illusion condition, two beeps and one flash were presented (2 beeps/1 flash). The SOA between the auditory stimuli varied between 50 ms, 100 ms, 150 ms, 200 ms, 250 ms, 300 ms, and 500 ms. Control conditions were also presented (2 beeps/2 flashes and 1 beep/1 flash), with control and 2 beeps/1 flash trials randomly permuted within a single block. Participants indicated the number of perceived flashes. In additional unimodal conditions, presented in separate blocks, participants indicated the number of beeps or flashes (see Suppl. Materials). Older adults perceived significantly more illusions than young adults [*F*(1,34) = 6.31, *partial eta*^*2*^ = 0.15, *p* = 0.02]. There was also a significant Age Group x SOA interaction in the 2 beeps/1 flash condition [F(6,204) = 3.19, *partial eta*^*2*^ = 0.09, p = 0.005], driven by more perceived illusions for older compared to younger adults between SOAs 100 ms – 500 ms (See Figure 1). Illusion perception between the age groups began to diverge at 100 ms (*p* = 0.002). These effects were not related to response bias, as there were no group differences in the multisensory control conditions (see Suppl. Materials, also for unimodal results).

**1.**
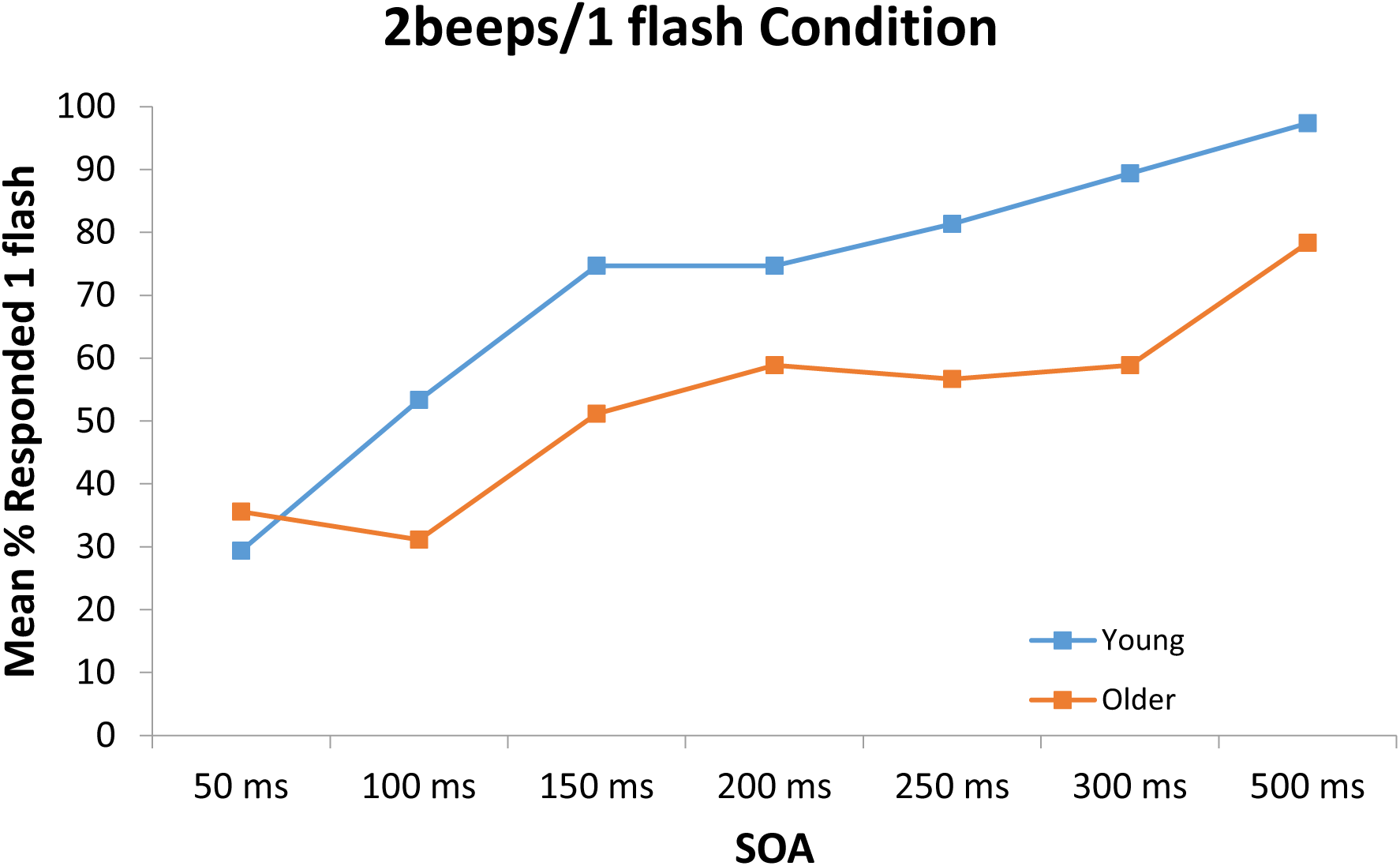
Behavioural results for the 2 beeps/1 flash condition. Older adults were less accurate (i.e., perceived more illusions) across the 100 ms – 500 ms SOA conditions, compared to young adults.

-----------Place Figure 1 about here -----------

### Neurophysiological signature of the illusory perception

In order to determine the neural activity underlying the increased illusion perception, both groups performed the same task inside the MEG -with the exception that only the 100-ms SOA: 2 beeps/1 flash condition, two-beeps, one-flash only, and two-flash conditions were presented. This was done to optimize the number of illusion trials. Epochs were cut to lengths of 1000 ms before the onset of the first audio-visual stimulus and 620 ms after the first stimulus.

To understand whether there was an age-independent factor determining a subject’s propensity for perceiving an illusion, we performed a 2x2 between-groups permutation-based ANOVA with factors Age (young vs. older) and Propensity for Illusions (perceived illusion (PPI) vs. perceived no illusion (PPNI); see Methods) on the squared amplitude of the sensor-level time-frequency transformed data. Special care was taken to define the appropriate permutations for a factorial design^14,15^ (see Methods). Older adults had significantly greater β-band activity (12Hz – 30Hz) compared to young adults in the time range of -250 ms – 75 ms relative to the onset of the first stimulus (*p* = 0.002). However, there was no main effect of the factor Illusion-propensity nor an interaction between both factors. There were no significant differences in other frequency bands.

We focussed our further network analysis of the MEG recordings to the time interval and frequency range of the differential sensory-level activity between the groups with a propensity to see the illusion vs. those with a propensity to see no illusion in order to determine how the network is used between groups. First, dynamic imaging of coherent sources (DICS)^16^, a frequency-domain adaptive spatial filtering algorithm, was used to identify the sources of the increased β-band activity. Beamforming revealed peak activity within this time interval to occur in the right middle temporal gyrus (50 -30 -10), right middle frontal gyrus (30 20 30), and bilateral fusiform gyrus (±20 -70 -10) (see Figure 2).

**2.**
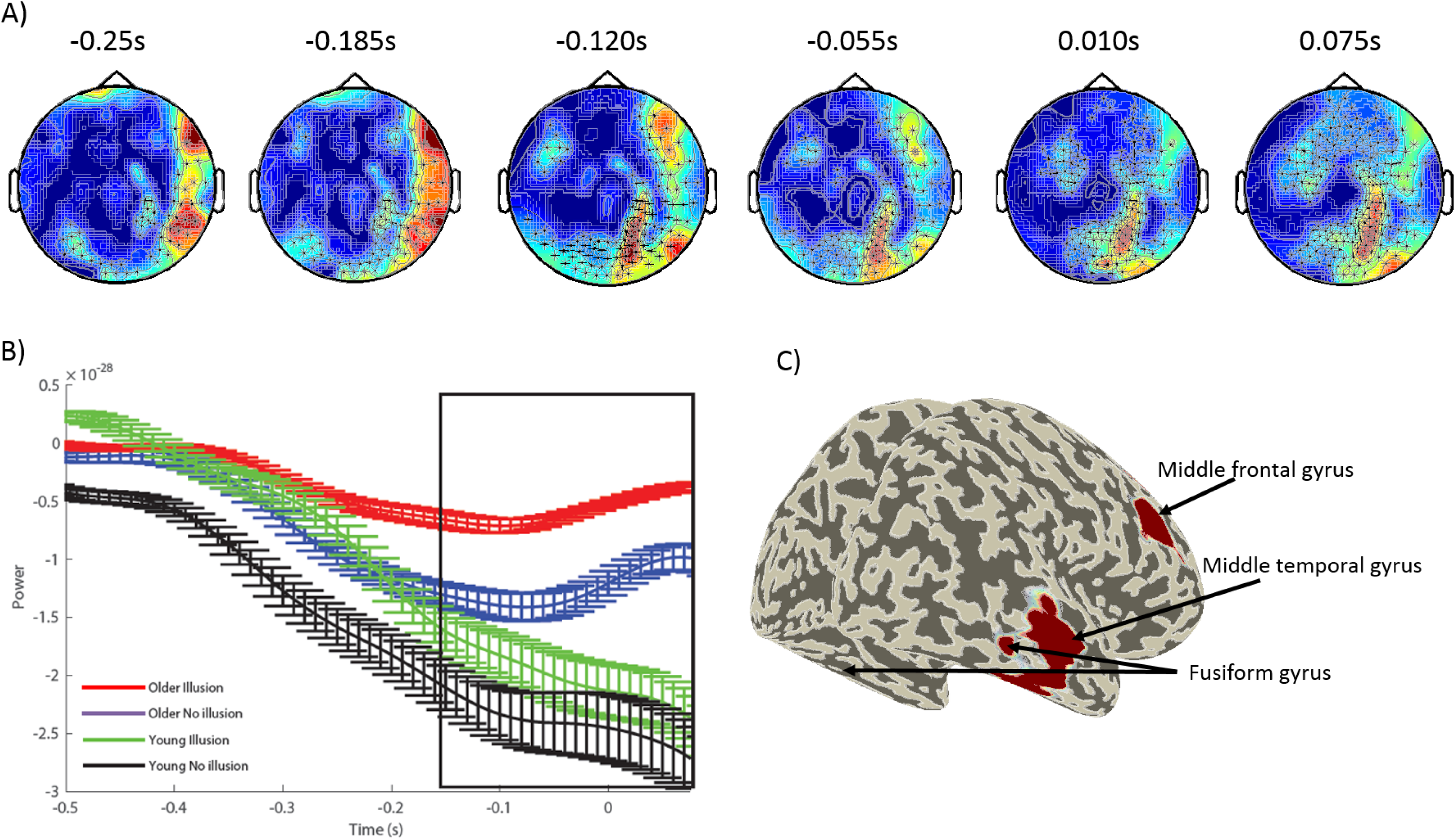
A) Topological plot representing the locations of the significant MEG channels for the main effect of Age Group. The crosses indicate the significant channels. B) The average β-band power over time for the significant channels. C) Source localisation results from the beamformer analysis.

-----------------Place Figure 2 about here -----------------

### Information Transfer within the Network

To infer the network underlying the perception of the SiFi, we used a combination of TE estimation and DCM. DCM uses Bayesian inference to obtain the most likely model of physiological interactions given the data. This Bayesian approach requires that plausible models enter the DCM analysis as priors. A common approach is to define models from relevant neural sources determined by beamforming. Furthermore, we were interested in the interaction between the sources of activity found here and the primary auditory (BA22) and visual areas (BA18). However, to explore the entire model space of the eight sources using DCM, would require the generation of 2^28^ = 268,435,456 models 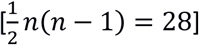 possible edges/connections] for *each* type of model connection (excitatory, inhibitory, and mixed).

This brute-force approach is computationally intractable. Furthermore, the models specified in DCM must be biophysically motivated and may not be randomly generated^14,15^. However, it is rarely the case that all effective connections (or lack thereof) are known between each of the cortical areas in question. To reduce the model space to a tractable size and to plausible models only, we took advantage of the MEG’s temporal precision and estimated transfer entropy (TE) between source time courses. TE is a model-free measure of information transfer between two processes^17^; the resulting network of information transfer between neural sources represents candidate connectivity relevant for solving a given task. Because we estimated bivariate TE from multiple sources it is very likely that some of the inferred links are indeed spurious due to cascade or common driver effects^18,19^. The TE network is thus a highly plausible starting point for building a model of effective connectivity underlying the neural computation but may be further refined by DCM.

We estimated delay-sensitive TE^20^ from reconstructed state spaces^21^ using individually optimized embedding parameters for each participant^22^ (see Suppl. Materials for details on TE estimation). We estimated TE for each pairwise combination of sources in each participant and tested these TE values for statistical significance using a permutation test against surrogate data^23^. We thus obtained networks for individual participants, which we then combined into group-level networks for propensity to perceived illusion (PPI) and propensity to perceived no illusion (PPNI), across the age groups, using a binomial test of individual links across all participants, irrespective of age (see Figure 3; see Suppl. Materials for a detailed description).

**3.**
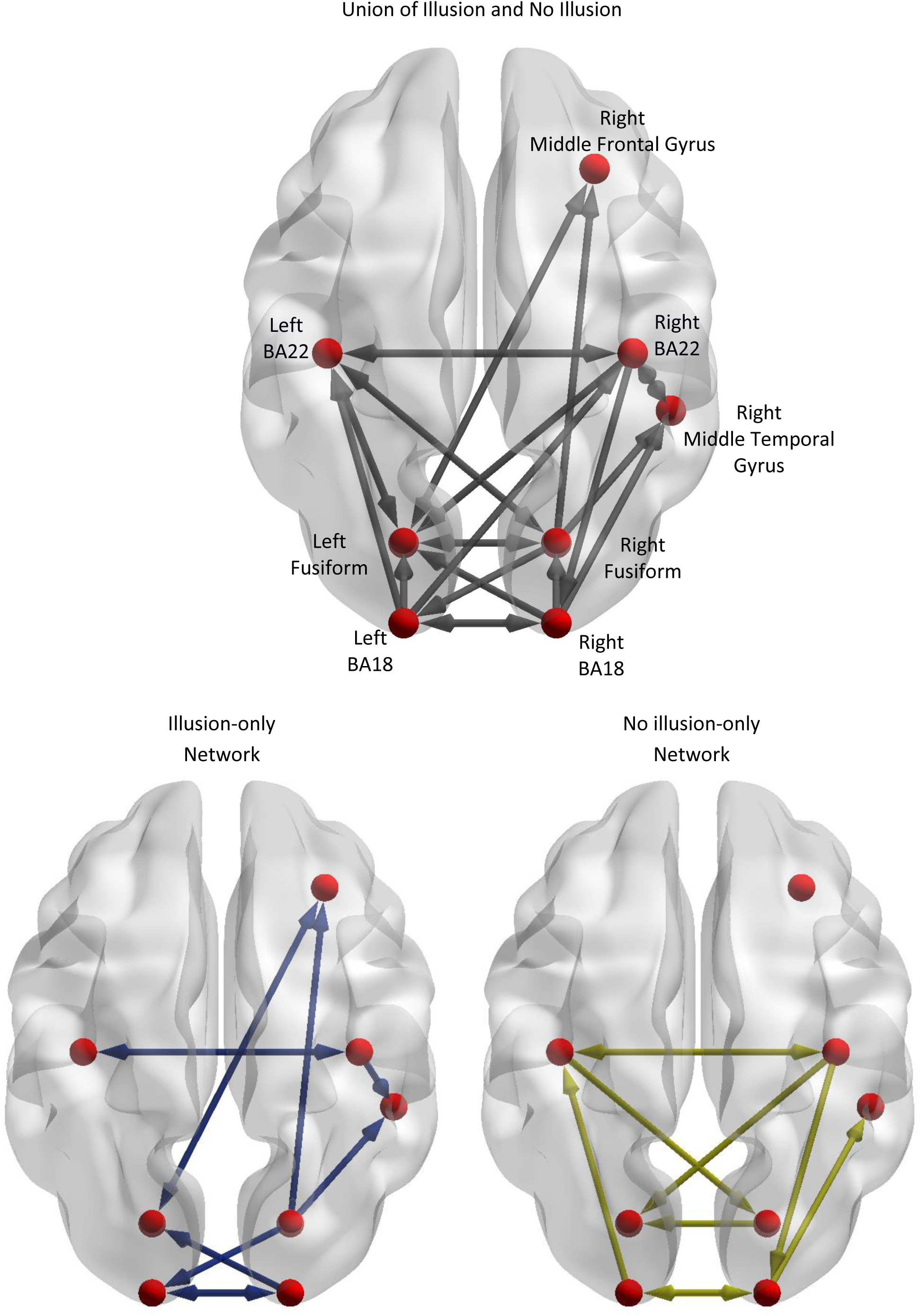
A) The combined results of the perceived illusion and perceived No illusion trials, from the transfer entropy analyses. This also includes links that are associated with both percepts. B) represents the differences between the perceived illusion-only and perceived no illusion-only conditions. Connections which are present in A) but not in B) indicate that those connections are present in both conditions.

-----------------Place Figure 3 about here -----------------

We then took the union of the group-level networks for the PPI and the PPNI groups and constructed DCM models to find the links that explained differences in performance between age groups. To avoid statistical ‘double-dipping’, for each participant TE estimation was performed on the odd-numbered trials and DCM on the remaining even trials. In order to confirm and refine the model, we used a systematic approach, where DCM was applied hierarchically in three steps, where the first two steps were aimed at obtaining a parsimonious, common model describing data for both groups of old and young participants while the third step then investigated age-related modulations of model parameters in this common model.

The aim of the first DCM analysis step was to determine if the union model or small variations of it offered a good description of data for age both groups and whether small variations to it would yield higher model evidence, indicating the need for a more thorough pruning of the union network. In this first TE inspired DCM analyses, for both age groups, twenty-four models were generated. The links for the DCM models were a union of the PPI and PPNI illusion TE models. The frequency range of interest was constrained to the β-band. Model 1 consisted of all links, with all links being active. Models 1.2-1.24 systematically removed one link, to identify any possible links whose removal might affect the model evidence. Models 1.25 and 1.26 consisted of only the links in either the illusion or no illusion TE models, respectively (see Figure 3). The resulting winning model was model 1.5, which was close to the union model, but had link Left BA18 to Right BA22 removed (see Table 1 in Suppl. Materials).

**Table 1.**
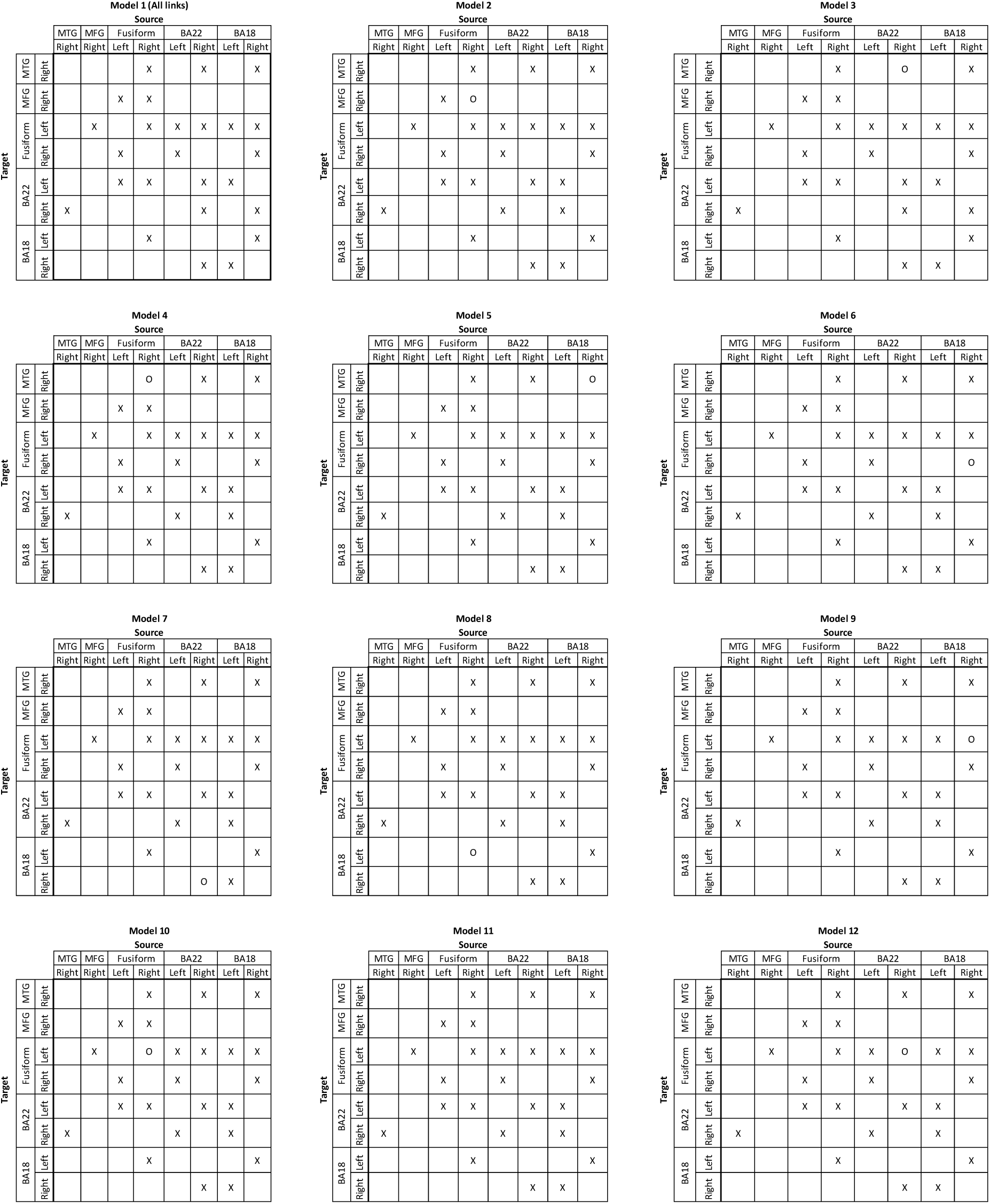
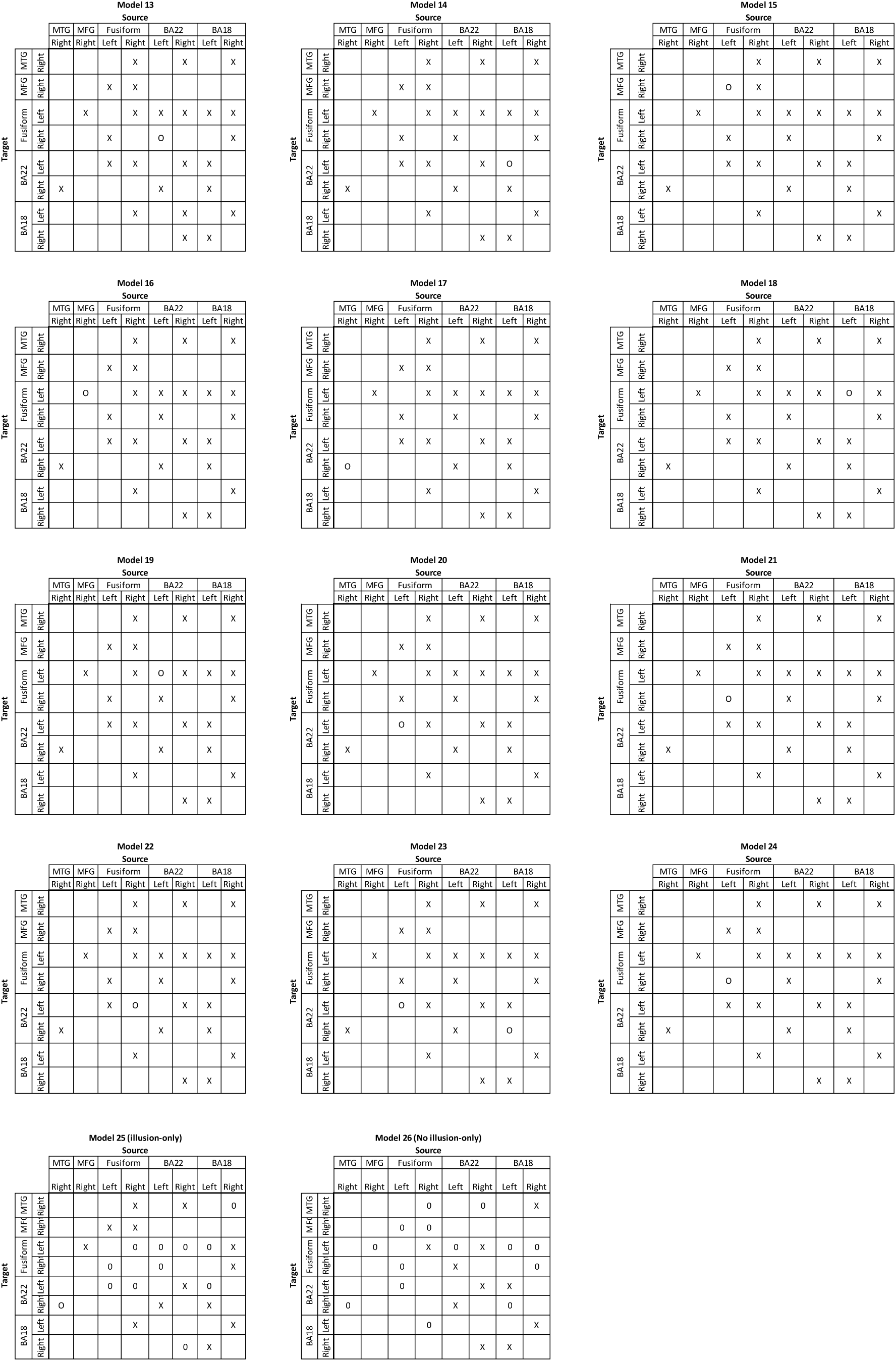
Tables of the links (X) which were included in each model. Model 1 contains all the links in the combined (illusion and no illusion) transfer entropy model. Models 23 and 24 represents the illusion-only and no illusion-only models from the transfer entropy analyses. For models 2-22, one link was removed. For the sake of clarity, “O” represent the missing links in each model. The winning model was model 9.

**Table 2.**
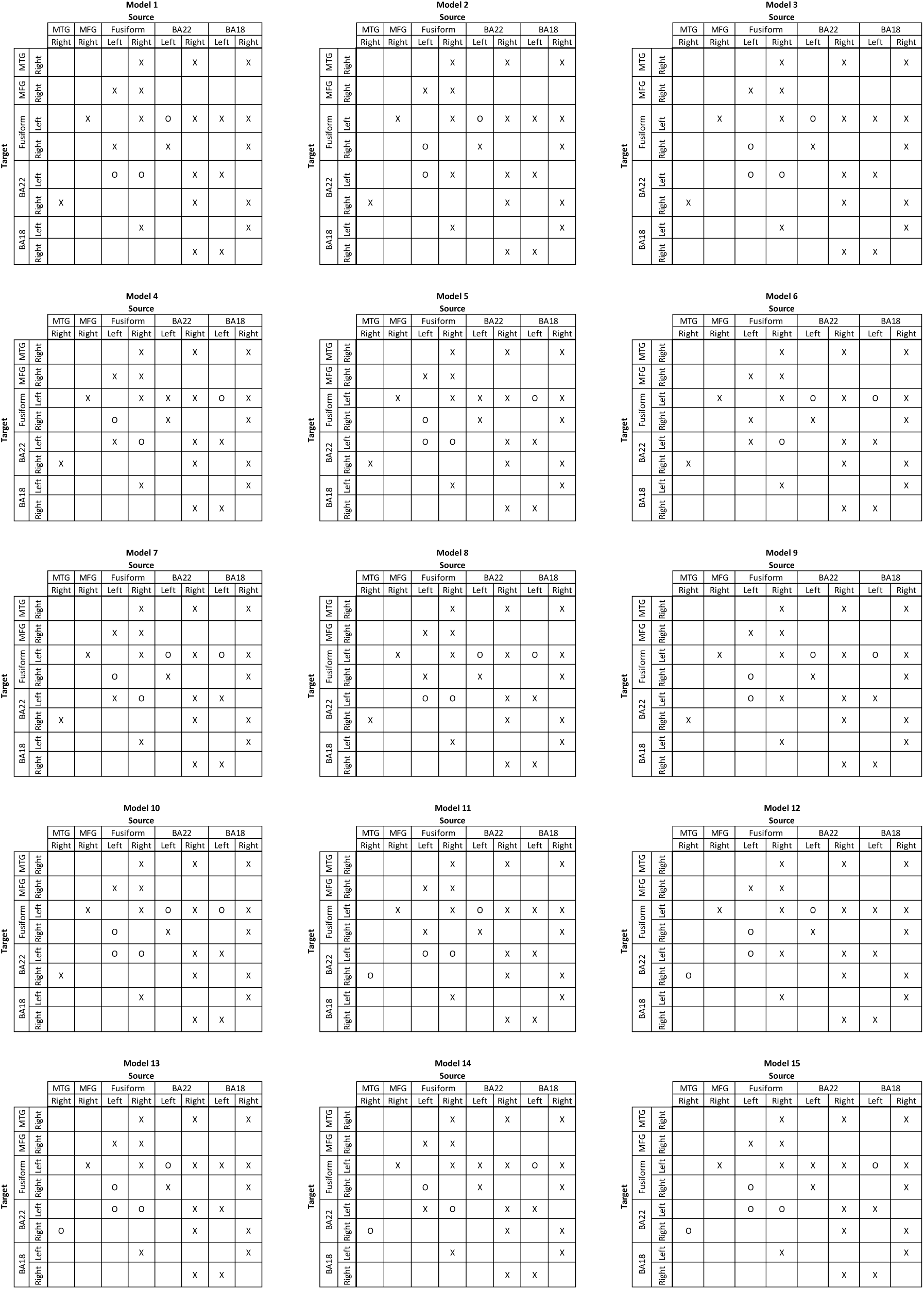
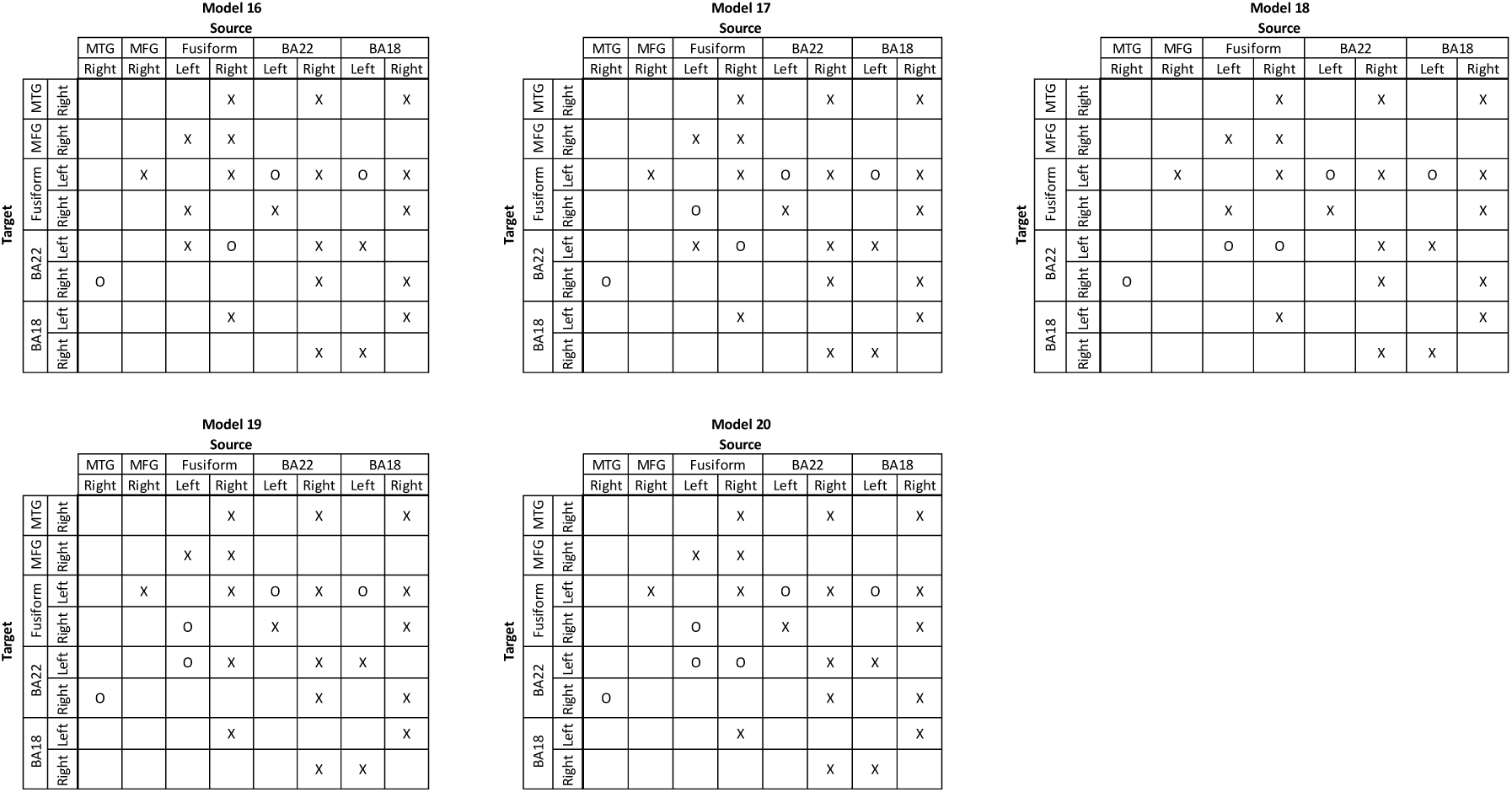
A separate set of DCM models were created to remove cascades and common drive artefacts. “X’s” represent the links in each model; “O’s” represent the *missing* links in each model. Models 2-20 are replications of models 1-10, respectively; with the exception that with each model, the link between the right middle temporal gyrus and right BA22 was also removed.

As the first DCM analysis step indicated an improvement of model evidence via pruning of links, a second DCM analysis was conducted to remove additional spurious effects due to common drives and cascade effects from the winning model (1.5) of the previous DCM analysis. This had to be done in a systematic and computationally tractable fashion^24-26^, because simply scanning models with all further possible combinations of pruned links would have resulted in a set of 2^27^ models. We therefore employed a strategy where pruned links constrained each other based on membership in acyclic triangles in the network graph (see Methods). The winning model for both the young and older adults was model 2.10 suggesting that the same network is used in both groups. We verified this by comparing the model distributions in the two groups via the model comparison statistics in the Variational Bayesian Analyses toolbox^27^. There was positive evidence against different model distributions between the two age groups (log Bayes factor = 6.68), confirming that both groups indeed employ the same neural network to perform the task (see Figure 4a).

**4.**
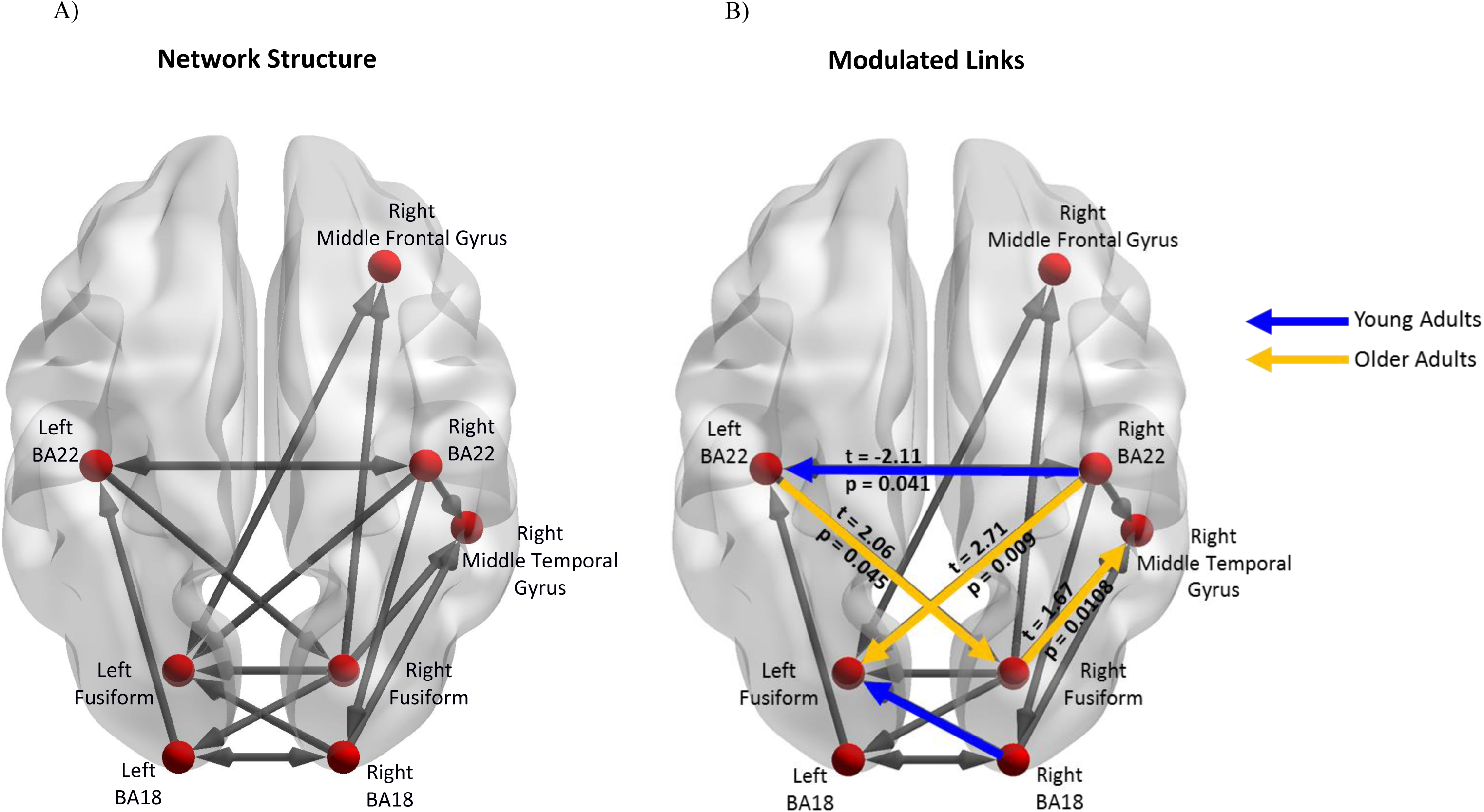
A) The winning model from the DCM analyses. This represents the structure of the network that is active in both young and older adults. B) represents the links which demonstrate significantly more β-band activity between the age groups.

The third stage of DCM analyses studied which links in the common network were modulated by the illusory percept in each age group. Thus, we examined the individual modulatory links in the binary connectivity matrix (*B*-matrix). The results from this DCM analysis found the winning model with respect to a modulation of connection strength by the illusory percept for the young adults was model 3.9 while the winning model for the older adults was model 3.7. Separate independent t-tests with factor age group were conducted for each modulation (see Figure 4b). Older adults showed increased modulation of links from: the left primary auditory (BA22) cortex to right fusiform cortex (*t* = 2.06, *p* = 0.045); right fusiform cortex to right middle temporal gyrus (*t* = 1.67, *p* = 0.011); and right auditory cortex (BA22) to left fusiform cortex (*t* = 2.71, *p* = 0.009), compared to the younger adults. Younger adults had greater modulation of the link from the right visual cortex (BA18) to the left fusiform cortex (*t* = -2.25, *p* = 0.03), and from right BA22 to left BA22 (*t* = -2.11, *p* = 0.041).

-----------------Place Figure 4 about here -----------------

## Discussion

Previous studies have demonstrated that older adults as well as some patient populations have increased rates of illusory percepts. Predictive coding theories offer a parsimonious explanation for this effect when taking into account that the amount of accumulated prior information increases over the life-span, while the precision of unisensory evidence decreases. Together these changes should favour interpretations of the world based on priors over veridical ones. Thus, comparing the neural basis of illusory percepts in young and old participants offers both, a critical test of microcircuit theories of predictive coding, as well as an opportunity for novel insights into perceptual changes in ageing.

We found that modulations of effective connectivity linked to illusory vs. veridical percepts carried clear signatures of cross-modal predictions in the older participants, whereas modulations in the younger participants were linked to changes in connections within unisensory networks only. This pattern of results aligns with a predictive coding account of ageing-related illusory perception. The link between the cross-modal use of priors and illusory percepts in older participants is further strengthened by the observation that older participants had increased activity in the β-band in the pre-stimulus phase. This strengthens the link because recent microcircuit theories of predictive coding suggest that priors or predictions are generated in cortical layers 5 and 6 and are signalled in the β-band^28^. A result seemingly conflicting with the link between β-band activity and priors (predictions) related to illusions is the fact that we found no significant effect of illusion-propensity across subjects on β-band activity. Yet, when analysing illusion vs. no-illusion conditions within subjects, we indeed found a trend towards increased pre-stimulus β-band activity for trials where an illusion is perceived compared to trials were no illusion is perceived (Suppl. Materials). Thus, we attribute the failure to observe β-band effects of illusion-propensity to the difficulty to separate age and illusion-related effects.

In addition to the more frequent occurrence of illusory precepts in older participants, they also exhibited more illusory percepts over a wider range of SOA. We interpret this as a widening of the acceptance window for binding auditory and visual events, brought about by the necessity to interpret degraded inputs from the visual system^4,29^.

We note that the illusion-related changes in effective connectivity as detected by our combined information-theoretic and DCM analysis were found between early and late sensory processing areas. This is in contrast to previous research that suggested that illusory percepts were caused by enhanced early sensory integration only, in young adults^30-32^. There are at least three reasons for this discrepancy between these findings. First, mechanisms related to cross-modal illusions may be detectable better in the older participant group as they are more dominant there, favouring their detection in our special study cohort. Second, transfer entropy compares favourably with other methods of finding connectivity^33,34^ as it is sensitive also to non-linear coupling (e.g. as required by communication between frequency bands). Third, the approach of model comparison based on DCM offers the possibility to disentangle effects due to structural differences in task-related networks (which were absent between groups) and illusion-related modulations of coupling strength (which were present and differed between groups).

Previous studies have suggested alternative explanations for illusory percepts in older people, e.g., relating them to an age-related delay in neural processing^35-37^. This is an unlikely explanation for our findings. When analysing the time course of the neural activity at the source-level, it was clear that the β-band activity in older adults was not delayed, compared to younger adults (see Figure 2B). In fact, for older adults who are more likely to perceive the illusion, the increase in amplitude of the β-band activity begins slightly earlier than their younger counterparts. Therefore, the results illustrated here are not likely due to neural delays caused by ageing.

From a purely methodological perspective, the current study is one of the first to combine exploratory (TE) and confirmatory (DCM) approaches to the analysis of network activity. While this has been suggested theoretically before for the combination of Granger Causality and DCM^13^, some comments related to the practical application are in order. First, we stress that applying the two analyses to separate data sets (e.g. odd and even numbered trials) is necessary to avoid variants of double dipping. Second, after deriving an estimate of the network structure from an exploratory approach, a confirmation of the network structure by model comparison requires the formulation of multiple plausible network models. Here, the original suggestion has been to use an increasing number of eigenmodes of the functional connectivity matrix for this purpose^38^. However, we here opted for a targeted removal of links from triangular network motifs indicative of common driver and cascade effects. This was done because such spurious links are known to appear in bivariate network analyses via transfer entropy, and were considered the main problem for the a priori validity of our models^25,39-41^.

In sum, our results suggest that the decrease of unisensory acuity and the accumulation of prior knowledge over the life span lead to a perception of the world increasingly dominated by this prior knowledge. Accordingly, older compared to younger adults had increased rates of illusory percepts, and showed modulations of cross-modal connections linked to these illusions. At the level of oscillatory neural activity both ageing and the behavioural occurrence of illusions were linked to increases in β-band activity. This supports recent neurophysiological accounts of predictive coding where priors and predictions are carried by β-band activity.

## Methods

### Participants

In this experiment, 25 healthy young adults (11 males) between the ages of 21-28 and 28 healthy older adults (12 males) between the ages of 58-72 took part. All participants were right-handed. Older adults were given the Consortium to Establish a Registry for Alzheimer disease (CERAD) questionnaire to ensure they did not suffer from age-related cognitive deficits^42,43^. All participants were also given the D2 attentional test^44^. All participants performed within their age-related norms.

Behavioural-only SiFi

### Apparatus and Stimuli

The visual stimuli were presented on a 24” flat panel computer monitor with a refresh rate of 60 Hz. The visual stimulus was a white circular disk, subtending 2° of visual angle. This disk was placed 4° of visual angle below the fixation cross. The presentation duration of the disk was 16 ms.

The auditory stimulus consisted of a 16 ms, 3500 Hz pure tone with a total rise- and decay-time of 20 μs and a sound pressure level at approximately 65 dBA. The auditory stimuli were presented using closed, circum-aural headphones (AKG, Austria, model: K271).

### Design and Procedures

The design of the experiment was based on a 3x7 repeated-measures design with Modality (vision-only, auditory-only, and audiovisual) and Stimulus-onset Asynchrony (SOA; 50 ms, 100 ms, 150 ms, 200 ms, 250 ms, 300 ms, and 500 ms) as factors. The dependent variable was accuracy. The factor Modality was blocked and the order randomized between participants. Participants received instructions and were given a short practice block to ensure they understood the task.

Within each block, the participants’ task was to indicate how many stimuli (visual or auditory) were presented. At the beginning of each trial, a fixation cross was presented at the centre of the computer screen. Participants were instructed to maintain their eye gaze on the cross throughout the experiment. If two stimuli were presented, the first stimulus was presented followed by a variable SOA, between 50 ms and 500 ms. Then, the second stimulus was presented. Afterwards, participants indicated via button press how many stimuli were presented. Each trial was followed by an inter-trial interval (ITI) between 1000 ms and 1500 ms (step sizes of 250 ms; see Figure 1). While reaction times were recorded, participants were asked to emphasize accuracy over speed. The experiment was programmed in Presentation (Neurobehavioral Systems, CA, USA).

----------------Place Figure 5 about here ----------------

**5.**
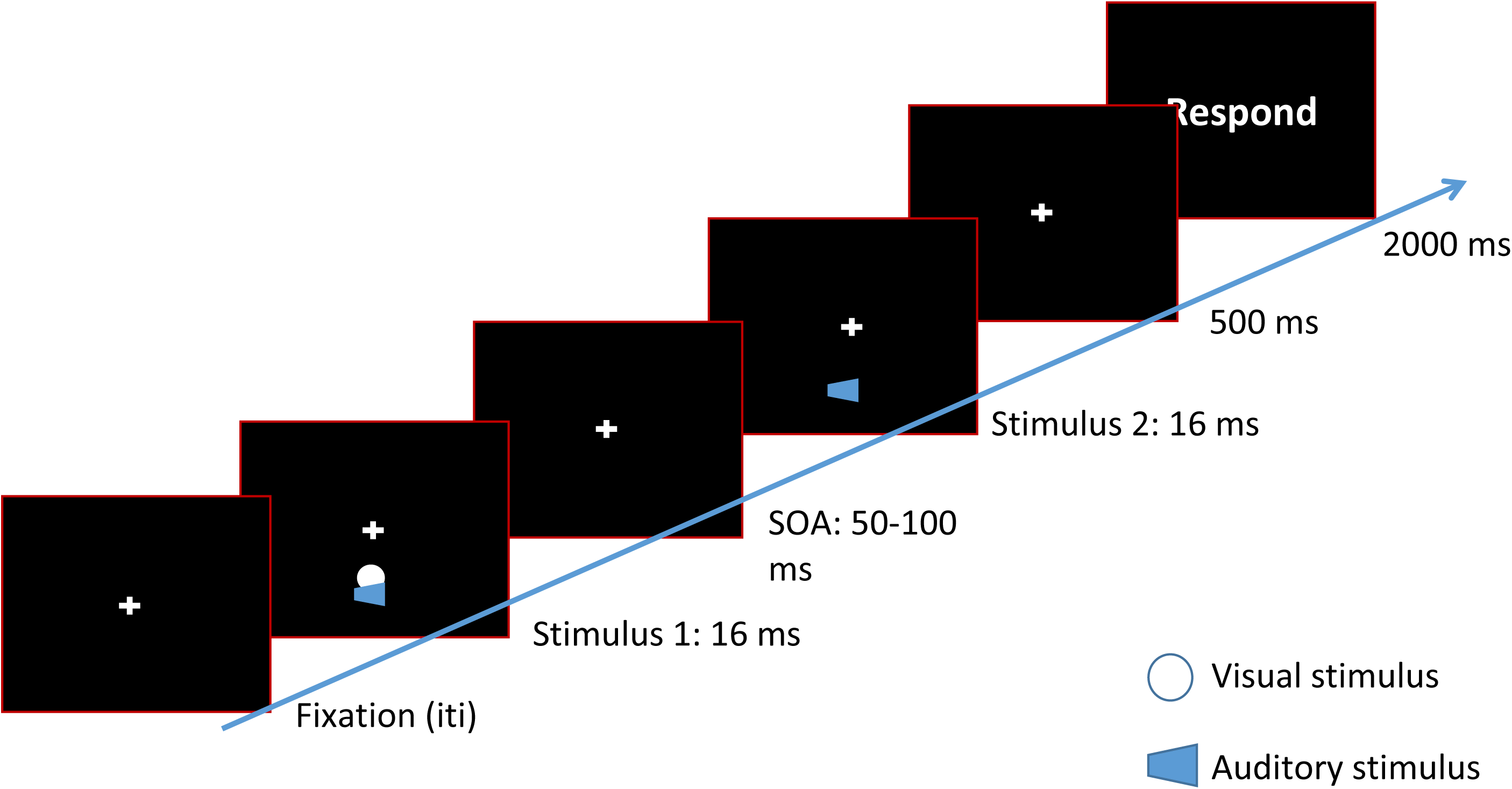
A timeline of a 2 beeps/1 flash condition. After a variable inter-trial interval, a white disk (flash) is presented along with a 3500 Hz tone (beep) for 16 ms. After a variable SOA, the second beep is presented. In the 2 beep/2 flashes condition a second white disk would be presented along with the second beep.

The unimodal-only blocks were separated into two separate blocks. In the vision-only block, one or two flashes were presented and the participant’s task was to indicate how many flashes were presented. In the auditory-only block, one or two beeps were presented and the participant’s task was to indicate how many beeps they heard. There were 140 trials in each of the unimodal blocks, 70 trials where one stimulus was presented and the remaining trials where two stimuli were presented, with an equal number divided between the SOA conditions.

The audiovisual block contained 210 trials in total. The audiovisual block contained two control conditions (1 beep/1 flash and 2 beeps/2 flashes) as well as the illusion condition (2 beeps/1 flash). In the illusion condition, the visual flash was presented at the same time as the first auditory beep. The second beep was presented at one of the previously indicated SOAs. In the control conditions, the audio-visual pairs were presented simultaneously. The second beep-flash pair in the 2 beep/2 flash control condition was presented at a variable SOA, relative to the first pair. This was done to minimize response bias towards responding to the auditory stimuli. The participants’ task was to ignore the auditory stimuli and indicate how many visual flashes were presented. All responses were made via computer keyboard.

MEG Methods

MEG was recorded in accordance to the suggested guidelines^45^. Participants were in a seated position, in a 275-channel, whole-head MEG with axial gradients (Omega 2005, VSM MedTech Ltd., BC, Canada). The sampling rate was 1200 Hz. Data were transformed to a synthetic third order axial gradient representation, and band-pass filtered in hardware between 0.1-300 Hz. Four EOG, two EMG, and two ECG electrodes were placed on the participant’s face and clavicle to record eye blinks, facial movements, and heart rate, respectively. Head localization was continuously recorded.

Visual stimuli were delivered via a video projector (Sanyo xp41) and back-projected to a semi-transparent screen at a distance of 60 cm from the participant’s head. Auditory stimuli were generated by a computer sound card (Creative Labs; Audigy 32) before going through sound conducting tubes into the MEG chamber. These sound-conducting tubes were connected to plastic ear moulds (ProPlugs, Doc’s). The sound pressure level was the same as in the behavioural experiment.

The task inside the MEG was similar to the task outside the scanner, with the following differences. There were four experimental conditions (2 beeps/1 flash, two-flashes, one-flash, and two-beeps). In the 2 beeps/1 flash, 2-beeps, and 2-flashes conditions, the second stimulus was always presented at an SOA of 100 ms. After 500 ms, the response screen was presented and participants indicated how many flashes were presented. In the two-beep condition, participants should respond ‘0 flashes’. Trials were rejected if participants responded before the response screen. The inter-trial interval was jittered between 850 ms – 1250 ms.

There were 200 repetitions in the 2 beeps/1 flash condition, 100 repetitions of single flash condition, and 100 repetitions of the two beeps condition. Fifty, 2-visual flashes trials were also presented. This was done to minimize the possible bias to respond ‘one flash’. Participants responded using a MEG compatible 5-button response box (Cambridge Research Systems, LTD) with their right hand. All trials were randomized across 5 experimental runs. Each run lasted approximately seven minutes. Participants received a short break between each run.

MEG analyses

### Pre-processing

All MEG data processing, except TE and DCM analyses, were carried out using FieldTrip^46^. Eight sensors (MRF22, MLT44, MRC12, MRC25, MRF22, MRO21, MRO53, and MRF11) had elevated noise levels and were thus excluded from the analyses.

Trials were cut from 1000 ms before the onset of the first audio-visual stimulus to 620 ms after the first stimulus. Trials containing eye blinks or muscle artefacts were removed by automated artefact rejection routines and subsequent visual inspection.

On average, young adults perceived 61.80% of 2 beep/1 flash trials (122 trials) as illusions, older adults perceived 64.74% of 2 beeps/1 flash trials (130 trials) as illusion. Three young and twelve older adults were removed from the analyses because of excessive artefacts. After artefact rejection, between 50–120 trials survived in each condition.

Based on their behavioural performance, participants within each age group were rank-ordered based on the proportion of perceived illusions. Then, a median-split was taken to place participants into two groups (Propensity to Perceive Illusion (PPI) and Propensity to Perceive No Illusion (PPNI)). As a result, after pre-processing, 12 young adults were placed in the PPI group and 10 young adults in the PPNI group. There were seven older adults placed in the PPI group, and nine older adults in the PPNI group (mean of perceived illusion in each group: Young PPI = 67.52%; Young PPNI = 11.57%; Older PPI = 10.44%; Older PPNI = 75.76%).

To ensure that the MEG results were not confounded by an uneven number of trials between the Perceived Illusion and Perceived No-Illusion conditions, stratified sampling was applied by randomly removing trials from the condition with the most trials until an equal number of trials was present in both conditions.

### Time-frequency analyses

A time-frequency analysis, using Morlet wavelets, was computed (wavelet length = 5 cycles, size of the Gaussian taper = 3). This procedure resulted in a single-trial estimation of 2 Hz to 60 Hz power, in 2 Hz steps. The baseline interval was between -1000 ms and -500 ms before the onset of the first stimulus. Finally, trials for each condition were averaged for each participant.

We extended the cluster-based permutation statistics implemented in the Fieldtrip toolbox to a 2x2 independent groups ANOVA, with factors Age (young vs. older) and Illusion (percentage of perceived illusion vs. perceived no illusion trials). Induced power for each condition was averaged within each participant then submitted to an ANOVA and F-values for the main effects or the interaction were computed for each sensor. Sensors where the F-value surpassed a critical F-value corresponding to an alpha level of 0.05 were selected and assigned to clusters based on their spatial adjacency. Neighbouring sensors were defined based on the template-approach implemented in Fieldtrip. The average minimum of neighbouring channels for the cluster analysis was 8.7 neighbours. Cluster-level statistics were calculated by taking the sum of the F-values within each cluster. These calculations were performed for each main effect and the interaction separately. The observed cluster-level statistics were then tested against the distribution of the maximum cluster-level statistics gained from Monte-Carlo simulations with 2000 permutations for each effect. At each permutation, group and condition assignments were shuffled and the estimation of F-values and the clustering procedure were repeated on the resampled data. The resulting maximum cluster values were used to construct the maximum cluster-level distribution under the null hypothesis. Clusters were considered to be significant at an alpha level of 0.05 if the originally observed cluster value was greater than the 95^th^ percentile of the maximum cluster-level statistic distribution. Cluster-based statistical tests effectively circumvent the multiple comparison problem by reducing the dependent variable to the maximum cluster size of neighbouring data bins showing the same effect^47^.

Special care has to be taken to define the appropriate permutations for a factorial design^48,49^. Permutations were restricted to occur within each factor (e.g. Age), while the assignment of participants to levels of the other factor (e.g. Illusion) was kept constant. For example, when testing the main effect of Age, the factor of propensity to perceive an illusion was held constant. No exact permutation tests based on the F-statistic exist for the interaction effect; since restricting permutation of the observations such that neither group main effect affects the corresponding F-ratio would leave no possible permutations of the data. An approximate test can be constructed by restricting permutations of factor levels to occur between one factor and subsequently permuting whole subjects across groups. Though variability due to the main effects is not held constant under such a permutation scheme, their variability impinges on all terms of the model, giving a reasonable approximate test.

### Source Reconstruction

Dynamic imaging of coherent sources (DICS), a frequency-domain adaptive spatial filtering algorithm was used to identify the sources of the effects found at the sensor-level, implemented in the Fieldtrip toolbox. While the DICS algorithm was designed to compute source coherence estimates, we used real-valued filter coefficients only and therefore restricted our analysis to the local source power^50^. The real part of the filters reflects the propagation of the magnetic fields from sources to sensors, as this process is supposed to happen instantaneously^51^. First, illusion and no-illusion trials were combined into one data set. Cross-spectral density matrices were computed for the task period of -250 – 0.75 ms, in β-band, based on the statistical analysis of spectral power at the sensor level (spectral smoothing indicated in parenthesis): 21 Hz (9 Hz). Subsequently, data from both response categories were projected separately into source space using the common spatial filter from the previous step. The source analysis was separately conducted on the activity of the two conditions, and the difference between the projected sources was tested for significance as described above. Source activity was interpolated onto individual anatomical images from magnetic resonance imaging (MRI) and subsequently normalized onto the standard Montreal Neurological Institute (MNI) brain using SPM8 in order to calculate group statistics and for illustrative purposes. A linearly constrained minimum variance (LCMV) approach was used to project the frequencies of interest into source space^24^. The combined filters were regularized at 5%^52^.

Beamformer filters were computed as “common filters” based on the activation and baseline data across all conditions. Using common filters for activation and baseline and all conditions allows for subsequent testing for differences between conditions; using common filters ensures that differences in source activity do not reflect differences between filters. Spatial filtering of the sensor data for source statistics was then performed by projecting single trials through the common filter for each condition separately.

### Connectivity Analyses

Previous studies using dynamic causal modelling (DCM) had to balance the number of models that needed to be created from identified sources versus computational time. This is because if all possible models were to be built from a set of identified sources and connections, this would result in an intractable model spaces as well as the inclusion of physiologically implausible models. To overcome this hurdle, we took advantage of the MEG’s temporal precision and performed transfer entropy analyses on half of the data (odd trials from all participants). Transfer entropy (TE) is a model-free measure of information transfer, it quantifies the additional information we can gain about a random process *Y* if we not only know *Y*’s past, but also the past of a second process *X*. Information transfer is then quantified as the conditional mutual information

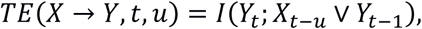

between the future value *Y*_*t*_ of the process *Y*at time *t*, and the past state *X*_*t*-*u*_, conditional on the past state*Y*_*t*-1_. Here, *u* is the reconstructed physical interaction delay *δ* between both processes. The delay is reconstructed by finding

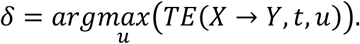

Prior to TE estimation, we reconstructed states *X*_*t*-*u*_, *Y*_*t*-1_ from scalar time series using a time-delay embedding^21^, with embedding parameters found through optimization of Ragwitz’ criterion^20^. Parameter optimization and delay-sensitive TE estimation from the ensemble of trials was done using the MATLAB toolbox TRENTOOL^23,41^, that implements the Kraskov-Stögbauer-Grassberger estimator for mutual information^53^. We used permutation testing against shuffled surrogate data to establish statistical significance for estimated TE values^23^.

We estimated TE for each possible pairwise connection in individual subjects, obtaining single-subject networks of information transfer. From single-subject networks we constructed group-level networks for the following groups: PPI and PPNI, by including links that were significant in at least 50% of the subjects within a group. The thresholding procedure corresponds to a one-sided Binomial test over subjects under the null hypothesis of significant links *k* being *B*(*k* V *p*_0_, *n*)-distributed, with *p*_0_ = 0.05 and *n* = 19 for the PPI group and *p*_0_ = 0.05 and *n* = 15 for the PPNI group. The threshold of 50% significant links is equivalent to an alpha level of 1e-10. We combined both group-level networks by taking the union of both sets of links.

Using the resulting TE network, the model space for DCM was dramatically reduced. DCM was performed using the Statistical Parametric Mapping, version 8 (SPM8), Matlab toolbox^53^. Overall, three separate DCM analyses were conducted. All DCMs were fit to the remaining (even) trials. As the activation occurred mostly before the presentation of the stimuli we employed a resting-state paradigm, using the linear neural mass model to calculate cross-spectral density of steady-state responses^54,55^. The time window was from 250 ms before stimuli onset to 75 ms post-stimuli onset. The data was detrended by simply removing the mean and the data was not subsampled. Eight modes were selected. Wavelet parameters were the same as those used to calculate the induced activity at the sensor-level.

A single equivalent current dipole for each source was selected as the electromagnetic model. The sources included into the model were the right middle temporal gyrus, right middle frontal gyrus, left fusiform gyrus, right fusiform gyrus, bilateral primary auditory cortex (Brodmann area 22), and bilateral primary visual cortex (Brodmann area 18). Network inputs were not selected given that this was a resting-state design. Random effects Bayesian model selection (BMS) was utilised to take into account the inter-individual variability in the structure of each model^56^. Separate model families were created for each type of interaction available (excitatory, inhibitory, and mixed). Then, a separate BMS was performed within the winning interaction-type family. In all cases, the family with purely excitatory interactions was the overall winner. Subsequently, the individual models within the excitatory family were compared using Bayesian Model Selection^57,58^.

For all DCM analysis steps, the union of the illusion and no illusion TE networks was the basis for investigated DCM models. A hierarchical approach was taken for the analyses, where the first two steps tested for spurious links in the TE network and the third step tested for modulations in the winning model. In the first DCM analysis step, Model 1.1 consisted of all links and all links were modulated. Models 1.2-1.24 consisted of all links, minus one to test for the possible presence spurious links that would trigger a second more in-depth analysis. All remaining links were modulated. Models and 1.26 consisted of only the links from TE models for the group with high illusion propensity and the group with low illusion propensity, respectively. The frequency range of interest for DCM was limited to the β-band, because the network nodes had been defined via the sources found in this band.

In the second DCM analysis step, triggered by the finding that pruning links increases model evidence, we tested for simple common drive and cascade effects by removing multiple potentially spurious network links simultaneously from the winning model from the first DCM step^25,26,40^, based on their membership in acyclic triangles in the directed network graph. Acyclic triangles may indicate spurious links, either due to one node driving the dynamics in the other two nodes (common drive effect), or due to a cascade of information transfer, where two consecutive links lead to spurious information transfer between the first and the third node (cascade effect)^18^. In such an acyclic triangle, only one of the two potentially spurious links can be actually spurious, because the two effects are mutually exclusive. This leads to a set of constraints on possible link-removals when trying to account for cascade and common driver effects.

Thus, in the alternative models, we systematically destroyed acyclic triangles, while making sure that no more than one of the two potentially spurious links was removed from all triangles in any given model. Possible combinations of simultaneously removable links were identified by encoding removable links as a Boolean function. This resulted in 20 possible models (see supplementary materials for a detailed list of all models).

After estimating model evidences for all candidate models, we tested the hypothesis that the model distributions differed between age groups versus the hypothesis that it was not different. To this end we used Bayesian group comparison as implemented in the Variational Bayesian Analyses toolbox^27^. Accordingly, the third step of the DCM analysis was carried out on the common winning model for both groups.

A third DCM analyses was conducted on the winning model from the second (refined) DCM analysis, to determine which links were modulated by illusory percepts. In this analysis, all links were maintained (A-matrix) but the modulation of individual links was systematically removed (B-matrix) (see Table 1 for a list of all β-band modulation values). We then statistically compared the illusion-trial related modulation in connectivity strength between the young and the older age group using t-tests.

## Supplementary Information

## Methods

### MRI Scanning Parameters

MRI images were collected using a 3 Tesla Siemens Allegra scanner (Siemens, Erlangen, Germany) at the Brain Imaging Center, Frankfurt using a 4 channel head coil. The subjects’ position was head-first supine and slice order was descending. A magnetization-prepared rapid acquisition gradient-echo (MP-RAGE) sequence was applied with the following parameters: TR: 2200 ms; TE: 3.93; flip angle: 9°; matrix: 256x256; FOV: 192 mm; voxel size: 1.0x1.0x1.0 mm3; number of slices: 160; distance factor: 50%. The duration of the sequence was 4 minutes. These scans were taken several weeks following the MEG testing.

### Results

### Behavioural

#### Unimodal

To determine the unimodal temporal acuity for the visual and auditory system alone, 2x2x8 Greenhouse-Geisser corrected mixed ANOVA was conducted with the factors Age Group (young vs. older) as the between-subjects factor, Modality (vision-only vs. auditory-only), and SOA (0 ms, 50 ms, 100 ms, 150 ms, 200 ms, 250 ms, 300 ms, and 500 ms) as the within-subjects factor. There was no main effect of Age Group [*F*(1,25) < 1, *n.s.*]. There was a main effect of Modality, with greater accuracy in the auditory-only condition (96.96%) compared to the vision-only condition (87.53%) [*F*(1,25) = 26.82, *p* < 0.0001]. There was also a main effect of SOA, with greater accuracy as the SOA increased (see Figure 1) [*F*(7,175) = 26.73, *p* < 0.0001]. There was a significant interaction between Modality and SOA [*F*(7,175) = 8.53, *p* < 0.0001]. A post-hoc analyses revealed that performance in the visual condition was significantly worse than the auditory condition in the 50 ms and 100 ms conditions (all *p*s = 0.05). Performance in the 50 ms SOA condition was significantly worse than all other SOAs in the vision-only condition (all *p*s = 0.05). There were no other significant interactions.

**Figure 1.**
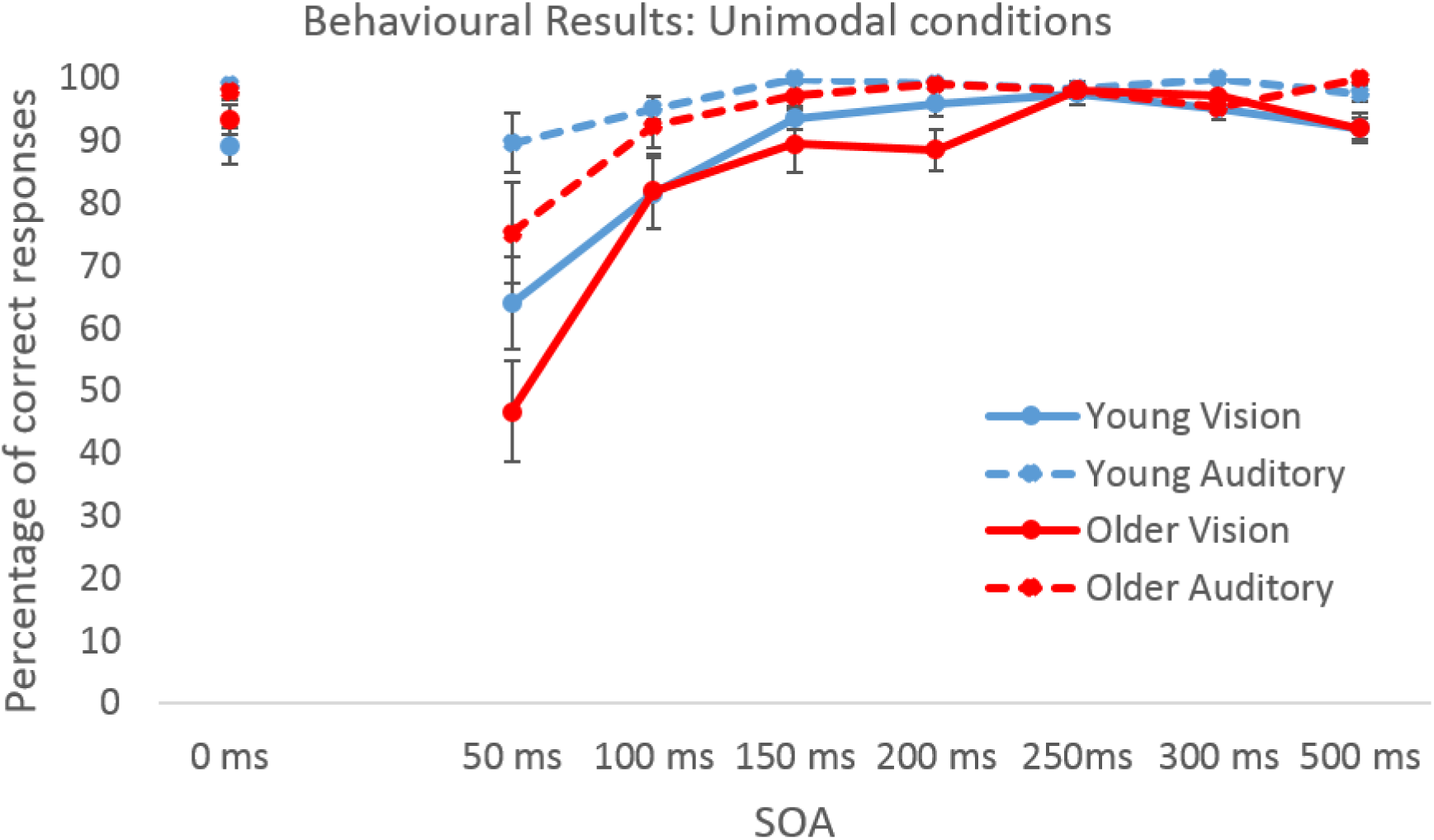
A graph of behavioural performance in the vision-only and auditory-only conditions. Overall, performance was better in the auditory-only conditions compared to the visual-only conditions. There was not significant difference between age groups.

#### Multisensory

To test for statistically relevant differences in accuracy, we conducted A 2x3x7 Greenhouse-Geisser corrected mixed-measures ANOVA was conducted with Age Group (young vs. older) as the between-subjects factor and Modality (2 flashes vs. 2 beeps vs. 2 beeps/1 flash) and SOA (50 ms, 100 ms, 150 ms, 200 ms, 250 ms, 300 ms, and 500 ms) as within-subjects factors.

There was a significant main effect between young (87.87%) and older adults (81.60%) [*F*(1,34) = 4.18, *p* = 0.05]. There was a significant main effect between the AV Conditions [*F*(2,68) = 54.10, *p* < 0.0001], with the highest accuracy in the 1 flash/1 beep (95.12%) and 2 beeps/2 flashes (95.63%) compared to the 2 beeps/1 flash condition (63.99%; all ps < 0.0001). There was a significant main effect of SOA [*F*(6,204) = 36.60, *p* < 0.0001], with accuracy improving as the SOA increased (50 ms = 71.24%; 100 ms = 78.46%; 150 ms = 84.98; 200 ms = 88.21%; 250 ms = 87.99%; 300 ms = 88.89%; 500 ms = 94.63%). There was a significant interaction between AV Condition and Age Group [*F*(2,68) = 5.36, *p* = 0.007]. A posthoc test revealed older adults were significantly worse in the 2 beeps/1 flash condition compared to other conditions and young adults, including performance from young adults in the 1 beeps/1 flash condition (all *ps* < 0.01). There was also a significant difference between young adults in the 2 beeps/1 flash compared to all other conditions (all *ps* < 0.01). Finally, there was a significant interaction between the three factors [*F*(12,408) = 3.20, *p* = 0.0002]. Another post hoc revealed performance in the 50 ms 2 beeps/1 flash condition was significantly worse when the older and younger participants (all *ps* < 0.05; see Figure 2). Older adults perceived more illusions compared to the younger adults, from 100ms-300ms (all *ps* < 0.05; see Figure 2). There were no significant differences between the two groups in the control conditions. Furthermore, there was no significant difference between the two groups in either unimodal conditions. The increase in perceived illusion for older adults was not related to a difference in unisensory acuity.

**Figure 2.**
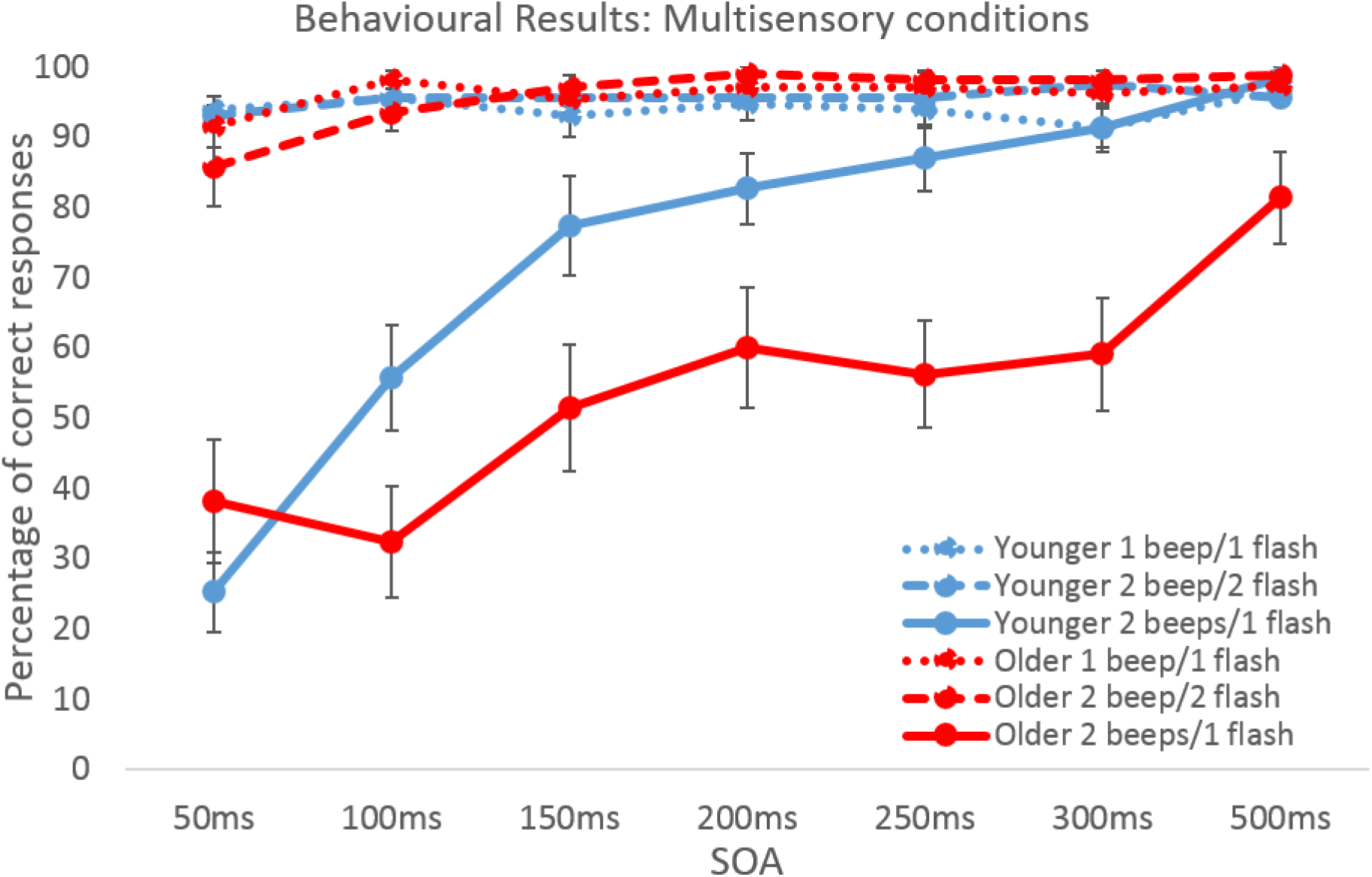
A graph of the multisensory conditions. There were no significant differences in the two control conditions (2 beeps/2 flashes and 2 beeps/1 flash) between the age groups.

#### MEG Results

A cluster-based permutation statistics, implemented in the Fieldtrip toolbox, to a 2x3 mixed-design ANOVA with Age Group (young vs. older) and the Illusion (illusion and no-illusion conditions) as the within participants sensor-level analysis revealed a trend towards increased β-band activity to trials where participants (regardless of age) perceived the 2-beeps/1flash condition as an illusion, compared to no illusion (*p* = 0.100; see Figure 3). This trend occurred between -5 ms – 9 ms over the frontal sensors then extending over the right parietal sensors.

**Figure 3.**
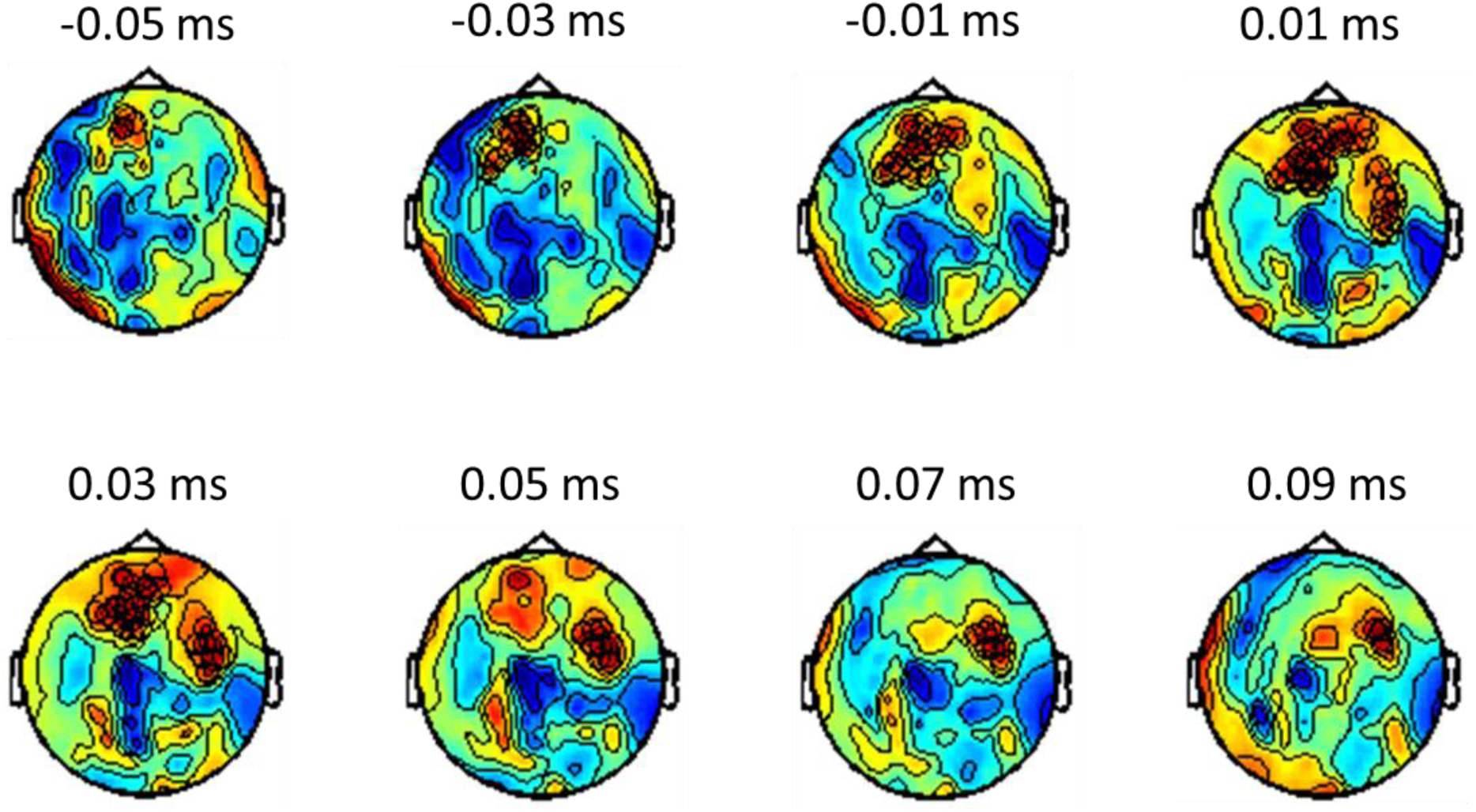
A figure representing the sensors associated with increased β-band activity in perceived illusion trials compared to perceived no-illusion trials. The sensors most associated with the increase in β-band activity are indicated by an “X”.

#### DCM results

The winning model for both the young and older adults was model 2.10 suggesting that the same network is used for both groups. To compare the Bayesian model Selection between the two groups, we used the model comparison statistics in the Variational Bayesian Analyses toolbox^1^. There was little model evidence for a different pattern of model evidence between each age group (model evidence = 6.6789; see Figure 1).

#### B0 results

The results from the third DCM analysis found the winning model for the young adults was model 3.9 while the winning model for the older adults was model 3.7. Separate independent t-tests were conducted for each modulation (see Figure 4 for a summary of results). Older adults showed increased modulation of the left primary auditory (BA22) cortex to right fusiform cortex (*t* = 2.06, *p* = 0.045); right fusiform cortex to right middle temporal gyrus (*t* = 1.67, *p* = 0.011); and right auditory cortex (BA22) to left fusiform cortex (*t* = 2.71, *p* = 0.009), compared to the younger adults. Younger adults had greater modulation from the right visual cortex (BA18) to the left fusiform cortex (*t* = -2.25, *p* = 0.03), and from right BA22 to left BA22 (*t* = -2.11, *p* = 0.041).

